# Recombinant B3 clade enterovirus D68 strains are efficiently rescued in 293T cells and infect human spinal cord organoids

**DOI:** 10.1101/2024.12.20.629498

**Authors:** Jennifer E. Jones, Sarah Maya, Gal Yovel, Jessica Ciomperlik-Patton, Jennifer Anstadt, Megan Culler Freeman

**Affiliations:** Department of Pediatrics, University of Pittsburgh School of Medicine, Pittsburgh, PA, USA; Department of Entomology, Texas A&M University, College Station, TX, USA; Center for Vaccine Innovation and Access, PATH, Seattle, WA, USA

**Author notes:** Address correspondence to Megan Culler Freeman,. Characteristics of recent recombinant EV-D68 strains.

## Abstract

Enterovirus D68 (EV-D68) is associated with respiratory disease in children. Between 2014 and 2018, biennial EV-D68 outbreaks coincided with peaks of a polio-like neurologic condition called acute flaccid myelitis (AFM). We hypothesized that specific mutations within the currently circulating B3 clade of EV-D68 impacted neurovirulence. However, recovery of these strains from infectious clones by published methods proved unsuccessful. Therefore, we tested different cell lines, reagents, and conditions to enhance efficiency of recombinant (r)EV-D68 rescue. In this study, we present a tractable rescue system to define virulence determinants in B3 clade strains. Using this approach, we successfully rescued historic and contemporary rEV-D68 strains to high titer after limited cell culture passage. All strains in our study replicated efficiently in the parental RD cell line and in human respiratory epithelial cell lines, with BEAS-2B cells exhibiting greater permissivity than A549 cells. While B2 and B3 clade strains could infect the SH-SY5Y neuroblastoma cell line, the neurovirulent B2 clade strain rUSA/IL/2014-18952 was more dependent on neuron differentiation than B3 clade strains and replication was not sustained under multi-cycle growth conditions. Conversely, replication of rUSA/IL/2014-18952 was more efficient in human spinal cord organoids, which model the cellular heterogeneity of the spinal cord, while replication of B3 clade strains was more modest. Immunofluorescence staining confirmed infection, with viral antigen colocalized with neurons. These findings suggest shifting dynamics of EV-D68 neurovirulence and provide a critical platform for further assessment of viral determinants of neurovirulence in B3 clade strains.

**IMPORTANCE:** Enterovirus D68 (EV-D68) can cause acute flaccid myelitis (AFM), a debilitating neurological condition of the spinal cord in children. Identifying viral determinants of EV- D68 neuropathogenesis is critical to understanding recent shifts in AFM prevalence; however, these investigations are limited to a small subset of infectious clones distantly related to currently circulating B3 clade strains. In this study, we leverage improved rescue strategies to characterize recombinant (r)EV-D68 strains from the dominant B3 clade. While all rEV-D68 replicate efficiently in the parental cell line and in human respiratory epithelial cells, B3 clade strains achieved greater titers in cultured neurons than a neurovirulent B2 clade strain, with less dependence on neuronal differentiation state. All B3 clade strains established infection in human spinal cord organoids, but replication varied between strains. Therefore, our study presents a tractable rescue system to begin to dissect viral determinants of shifting neurotropism within contemporary EV-D68 clades.

## INTRODUCTION

Enterovirus D68 (EV-D68) is a non-enveloped member of the human enteroviruses in the *Picornaviridae* family (1). EV-D68 was first discovered in 1962 in oropharyngeal swabs from four children with pneumonia and bronchiolitis; one of these prototype strains is named ‘Fermon’ (1). EV-D68 cases were rarely identified until 2005, when outbreaks of severe respiratory tract infections of EV-D68 were reported worldwide (2–4). This corresponded with viral diversification into three primary clades: A, B, and C (3–8).

EV-D68 has been associated with acute flaccid myelitis (AFM), a polio-like neurologic condition causing paralysis primarily in children, since 2014 (9, 10), though it was identified in a case report as early as 2008 (11). Due to this, EV-D68 is now considered a reemerging pathogen. In parallel with its novel association with AFM, EV-D68 continued to diversify into clades A1, A2, B1, B2, and B3 (10, 12–16). Biennial outbreaks of EV-D68-associated AFM recurred between 2014 and 2018 until viral transmission was disrupted in 2020 by nonpharmaceutical interventions implemented during the COVID-19 pandemic (17–23). EV-D68 circulation resumed in 2022, but was not associated with outbreaks of AFM for the first time since 2014 (23, 24). At the same time, the B3 clade grew increasingly dominant, with all but one of the sequenced isolates from 2022 falling in this clade and continuing to diversify into a new cluster (nextstrain.org) (25–27). Whether and how this new wave of diversification contributed to the stark decline in AFM prevalence is unclear.

Investigation into clade-specific determinants of EV-D68 neuropathogenesis requires a robust reverse genetics system. Infectious clones for EV-D68 were first engineered from the Fermon strain in 2018 by two research groups in China (28, 29). Sun *et. al.* cloned the viral genome under a T7 promoter, whereas Pan *et. al.* used a human RNA Pol I promoter. In both studies, recombinant (r) Fermon virus was successfully recovered after transfection of 293T or RD cells directly with infectious clone DNA (with co-transfection of T7 polymerase in the case of Sun *et. al.*), but Pan *et. al.* noted a PCR- induced mutation in the 5’UTR that impaired replication (28, 29). Sun and colleagues later went on to apply their reverse genetics method to generate an infectious clone from a neurovirulent B2 clade strain (30). Two years later, infectious clones for Fermon as well as a limited number of B1 and B2 clade strains were generated in the US under a T7 promoter and deposited to the Biodefense and Emerging Infections Research Resources Repository (BEI Resources), making these the first widely available infectious clones for EV-D68 (31). Rather than transfection, the authors recovered infectious virus by electroporation of *in vitro* transcribed viral RNA into RD cells (31). To date, no published infectious clones exist for B3 clade strains, but additional A1 and B3 clade infectious clones under a T7 promoter have since been deposited to BEI by the CDC. We and others were unable to recover infectious virus from these plasmids by electroporation of RD cells. Therefore, despite the growing availability of EV-D68 infectious clones, the study of viral determinants of EV-D68 neuropathogenesis remains restricted to strains that are increasingly genetically distant from those currently circulating.

In this study, we set out to establish a robust rescue platform of rEV-D68 from publicly available infectious clones. We compare rescue of rEV-D68 in BSR-T7, a sub-clone of BHK-21 cells that constitutively express T7 polymerase (32), RD, and 293T cells. We demonstrate that while rEV-D68 rescue is achievable in all cell types, rescue efficiency is most effective in 293T cells with T7 polymerase co-transfected in excess of infectious clone. This approach was well-suited to rescue of diverse rEV-D68 strains, including the prototype Fermon and viruses from A1, B2, and B3 clades. We further report that while all rEV-D68 replicate efficiently in RD cells and in human respiratory epithelial cells, only viruses of the B2 and B3 clades can infect human neuroblastoma cells. Finally, we determine that human spinal cord organoids are susceptible to infection with recombinant B3 clade viruses. Altogether, we present a reproducible system for rescue of contemporary neurotropic B3 clade viruses.

## RESULTS

### Rescue efficiency of recombinant (r)EV-D68 is impacted by cell type and ratio of polymerase to infectious clone

Discrepancies between research groups in choice of cell line and conditions for delivery of viral genome have created ambiguity surrounding ideal rescue conditions for rEV- D68 (28, 29, 31). Therefore, we began with a comprehensive investigation of rEV-D68 rescue in three candidate cell lines: BSR-T7, RD, and 293T cells. To establish optimal rEV-D68 rescue conditions, we first defined the transfection efficiency of each cell type. We optimized transfection efficiency in each cell line by titrating the concentration of a GFP expression plasmid, pcDNA 3.1 (+) CT-GFP, with either Lipofectamine 3000 or TransIT-LT1 (**Figure 1A**). We scaled up transfection conditions for rescue of rEV-D68 in each cell line with a well-characterized strain, rUSA/IL/2014-18952, since it is already readily rescued by published means (31, 33). We tested whether polymerase availability in RDs and 293Ts impacts rescue efficiency using different ratios of T7 polymerase to infectious clone. The recovered virus readily induced observable CPE in transfected BSR-T7 and 293T cells by 48h and in RD cells by 72h. Viral stocks were titered directly after transfection, to assess viral yield without additional rounds of passage. While a recent study reported that infectious rEV-D68 is not readily detectable in 293T cells after transfection (34), we found that all three cell lines produced infectious virus in all conditions tested (**Figure 1B**). Moreover, rEV-D68 rescue was most efficient in 293T cells, with titers 1 to 3 logs higher than RD or BSR-T7 cells. Viral titers were highest in 293T cells, consistently exceeding 10^6^ PFU per ml, particularly at the highest ratio of polymerase to infectious clone. In contrast, the lowest viral titers (10^4^ to 10^5^ PFU per ml) were observed in RD cells, especially at the lowest ratios of polymerase to infectious clone. The ratio of polymerase to infectious clone was more strongly correlated with infectious virus titer in RD cells than in 293T cells (adjusted R^2^ = 0.7 and 0.4, respectively). Thus, we concluded that rescue of rEV-D68 is most efficient in 293T cells at high ratios of polymerase to infectious clone.

**Figure 1.**
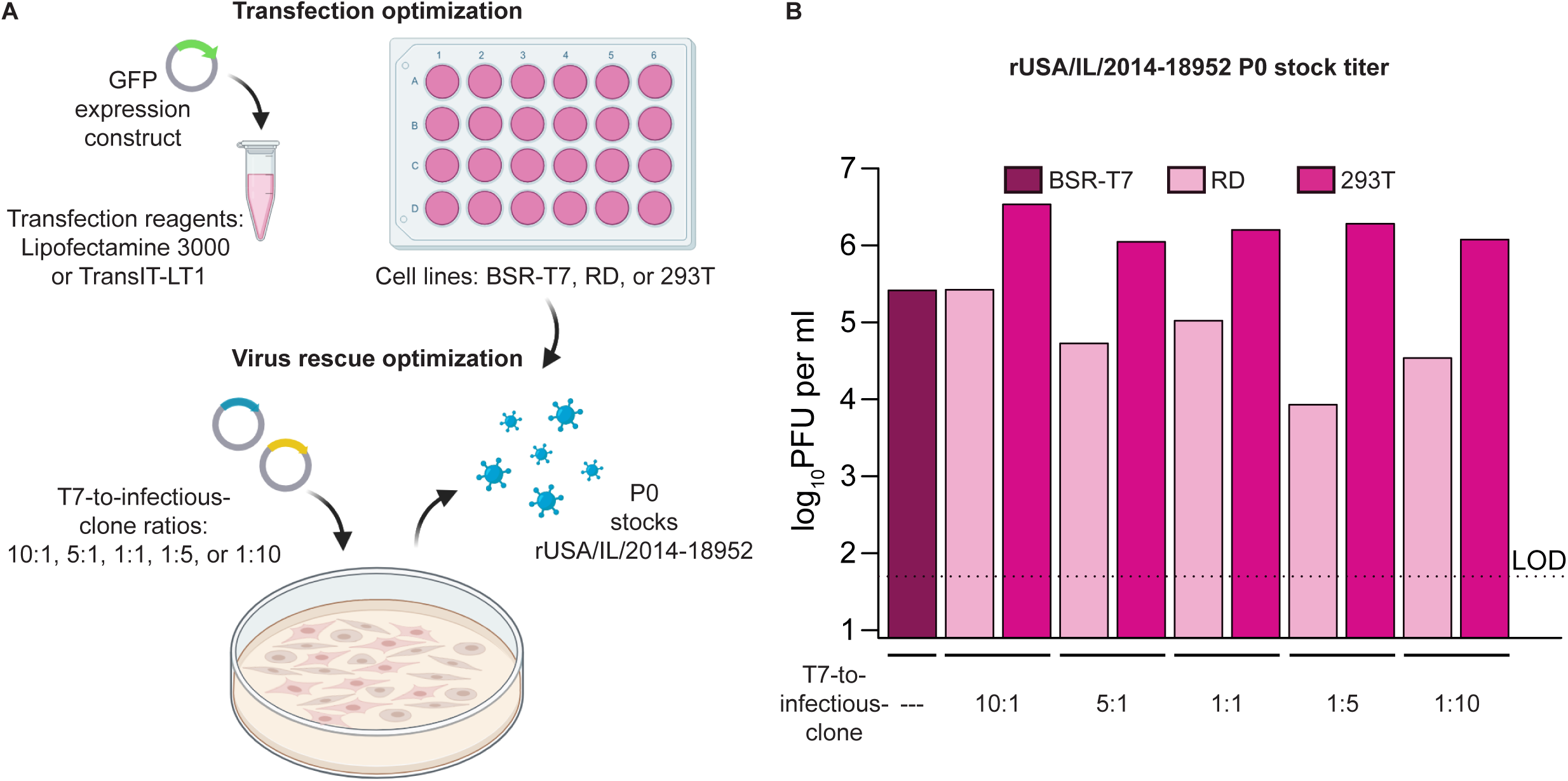
Rescue of rUSA/IL/2014-18952 from infectious clone is most efficient in 293T cells. **(A)** Infectious clone rescue optimization schema. Transfection conditions were first optimized in each cell line with a GFP expression construct. Optimal transfection conditions were applied to virus rescue. BSR-T7, RD, or 293T cells were transfected with pUC19-EVD68_49131. RD and 293T cells were co-transfected with pCAGGS T7 at a ratio of 10:1, 5:1, 1:1, 1:5, or 1:10. Mock-transfected cells received transfection mixture without DNA. Cells were incubated at 33°C, 5% CO_2_. Cells were scraped from the dish and collected with supernatant (designated P0 stocks). Created with Biorender.com. **(B)** Viral titers were quantified in P0 stocks from each condition of Figure 1A by plaque assay in RD cells.

### Recombinant B3 clade strains are efficiently rescued in 293T cells

Rescue of rEV-D68 from some previously circulating clades is well-established (28–31, 33, 34), but to our knowledge, infectious clones for B3 clade strains remain unpublished despite their availability through BEI Resources. Therefore, we assessed whether the method we optimized for rUSA/IL/2014-18952, a B2 clade strain, facilitated rescue of virus from these infectious clones. Infectious clones were available for USA/FL/2016- 19504, USA/OH/2018-23088, and USA/IL/2018-23252. Two additional infectious clones were available through BEI Resources for strains isolated prior to the association of EV- D68 with AFM in 2014: USA/Fermon (infectious clone deposited by (31)) and USA/WI/2009-23230. These strains were included in this study as controls as they were not expected to be neurotropic (35, 36). To confirm clade assignments, we reconstructed a phylogenetic tree of VP1 sequences from all five of these strains as well as the neurovirulent B2 clade strain USA/IL/2014-18952. As expected, sequences from B3 clade strains USA/FL/2016-19504, USA/OH/2018-23088, and USA/IL/2018-23252 formed a cluster distinct from USA/IL/2014-18952 (**Figure 2A**). Strains that preceded the 2014 association of EV-D68 with AFM, USA/Fermon and USA/WI/2009-23230, were distantly related to B2 and B3 clade strains.

**Figure 2.**
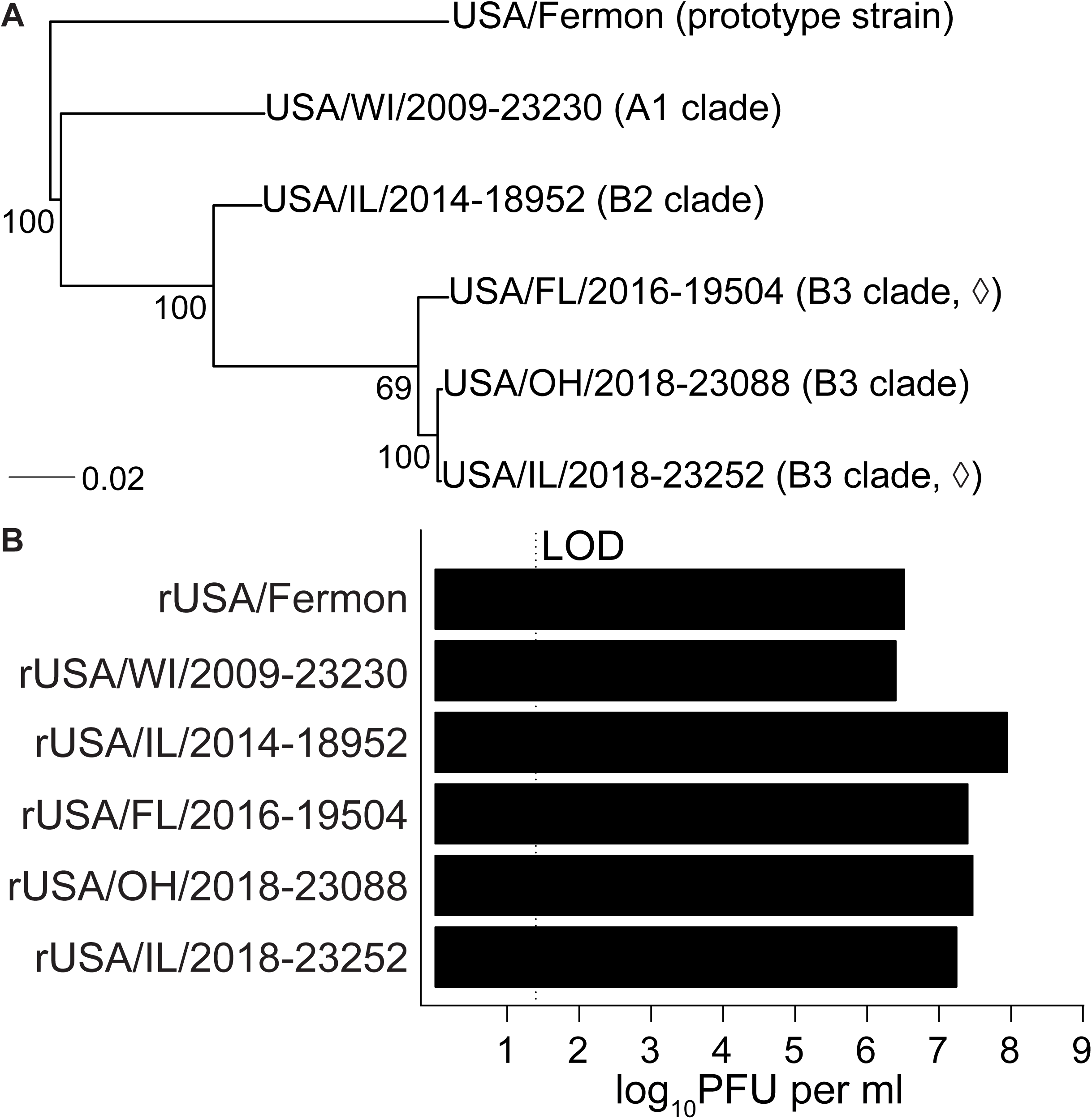
Contemporary EV-D68 infectious clones are efficiently rescued in 293T cells. **(A)** A maximum likelihood phylogenetic tree was reconstructed from VP1 sequences obtained from GenBank. Bootstrap values are for 1,000 trees. Scale bar indicates substitutions per site. (◊) denotes confirmed AFM. **(B)** A full panel of rEV-D68 strains was rescued in 293T cells and amplified on RD cells. Viral titers in P1 stocks were quantified by plaque assay in RD cells.

We next tested our rescue strategy in all six strains. Recently, Choi *et. al.* determined that enterovirus replication is sub-optimal during rescue in 293T cells (34); thus, we amplified virus in one additional passage (P1) in RD cells. All six rEV-D68 produced infectious virus from P1 stocks at titers of 10^6^ PFU per ml or higher (**Figure 2B**). Titers for rUSA/IL/2014-18952 were highest, reaching 10^8^ PFU per ml. B3 clade strain titers exceeded 10^7^ PFU per ml and were higher than those for control strains. Thus, our method permits efficient rescue of B3 clade rEV-D68.

### Recombinant B3 clade strains replicate efficiently in the parental RD cell rescue line and human respiratory epithelial cell lines

At present, few studies have investigated cellular tropism of B3 clade EV-D68 isolates (36, 37), and none used recombinant viruses. Therefore, we established the replication kinetics of this panel of rEV-D68 in susceptible cell types, beginning with the parental RD cell line on which our viral stocks were passaged. As expected, all strains replicated efficiently in RD cells under single- and multi-cycle growth conditions (**Figure 3, green circles**). Thus, our rescue method consistently produces replication-competent rEV- D68.

**Figure 3.**
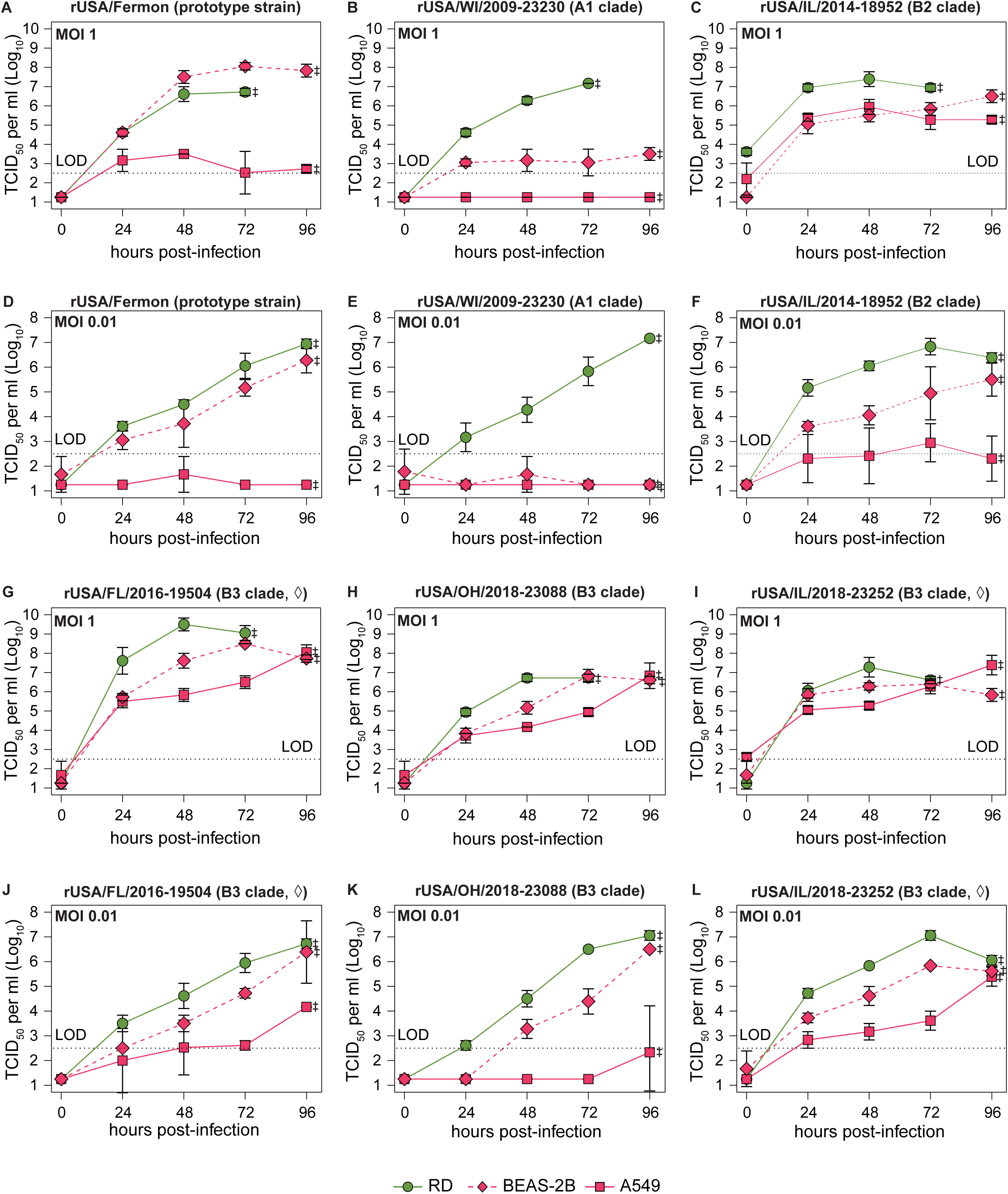
rEV-D68 strains replicate efficiently in parental RD and human respiratory epithelial cell lines. RD, BEAS-2B, and A549 cells were infected in triplicate with the indicated rEV-D68 strains at an MOI of 1 **(A-C** and **G-I)** or 0.01 **(D-F** and **J-L)**. Supernatants were collected at the indicated time points. Endpoint titers represent combined cell lysate and supernatant, as denoted by (‡). (◊) denotes confirmed AFM. Viral titers were quantified by TCID_50_ assay on RD cells. Error bars indicate standard deviation.

Since EV-D68 is most often detected in upper respiratory tract specimens (10, 38), we expected all of the rescued viruses to replicate efficiently in respiratory cells. Therefore, we examined replication kinetics in two human respiratory epithelial cell lines: BEAS-2B and A549. Each recombinant B3 clade virus replicated efficiently in these cells at an MOI of 1 (**Figure 3G-I, diamonds or squares, respectively**). Interestingly, all three B3 clade strains reached greater titers in BEAS-2B cells over A549 cells at an MOI of 0.01 (**Figure 3J-L, diamonds vs. squares**). Similar results were observed for the B2 clade strain rUSA/IL/2014-18952 (**Figure 3C** and **3F, diamonds and squares**). Differences between BEAS-2Bs and A549s were even more pronounced in control strains, rFermon and rUSA/WI/2009-23230, which replicated poorly in A549 cells even at high MOI (**Figure 3A-B** and **3D-E, diamonds and squares**). In fact, the only strain that exhibited robust replication in A549 cells at an MOI of 0.01 was the B3 clade strain rUSA/IL/2018- 23252 (**Figure 3L, squares**). The A1 clade strain rUSA/WI/2009-23230 was the only strain unable to replicate to high titer in either cell type at either MOI, though replication in RD cells was preserved (**Figure 3B** and **3E**). Overall, these data indicate that recombinant B3 clade viruses replicate efficiently in respiratory epithelial cells in addition to the parental RD cell line, but replicative differences exist between human respiratory cell lines that are not clade-specific.

### Recombinant B3 clade viruses efficiently infect neurons

EV-D68 has been associated with large outbreaks of AFM since 2014 (9, 10). Some, but not all, rEV-D68 from previously circulating B1 and B2 clades can invade the CNS in mouse models (30, 31, 33, 39), but rEV-D68 from the currently circulating B3 clade have not been evaluated for neurotropism. Therefore, we examined whether our panel of recombinant B3 clade strains are neurotropic in cell culture models. We first explored replication kinetics of our B3 clade viruses in the human neuroblastoma SH-SY5Y cell line, an established model for EV-D68 neurotropism (35, 37). Infectious virus was not reliably detected in SH-SY5Y cells infected with control rEV-D68 strains at the MOIs tested (**Figure 4A-B** and **4D-E**), consistent with previous literature suggesting that neurotropism in EV-D68 is not observed in strains isolated prior to 2014 (9, 10, 35–38). Infectious virus was detectable at 24 hours post-infection (hpi) at an MOI of 1 for the B2 clade rUSA/IL/2014-18952 strain, although replication was abortive (**Figure 4C, solid line**). In contrast, all B3 clade viruses replicated efficiently in SH-SY5Y cells at an MOI of 1 (**Figure 4G-I, solid lines**). Surprisingly, each B3 clade strain achieved a peak titer of 10^5^ to 10^6^ TCID_50_ per ml, several logs higher than the neurotropic rUSA/IL/2014- 18952 strain at the same MOI (**Figure 4C, solid line**). At an MOI of 0.01, differences between neurotropic strains emerged. Infectious virus was not consistently detected for the B2 clade rUSA/IL/2014-18952 strain at this MOI (**Figure 4F, solid line**). Among B3 clade strains, rUSA/OH/2018-23088 only produced detectable virus at the experimental endpoint of 96 hpi and only when cells were differentiated (**Figure 4K**). Likewise, replication of rUSA/FL/2016-19504 was markedly delayed (**Figure 4J**). Only rUSA/IL/2018-23252 exhibited sustained logarithmic growth in these cells at this MOI (**Figure 4L**). These data demonstrate that B3 clade rEV-D68 strains efficiently infect cultured neurons but differ in their capacity for continuous replication under low MOI conditions.

**Figure 4.**
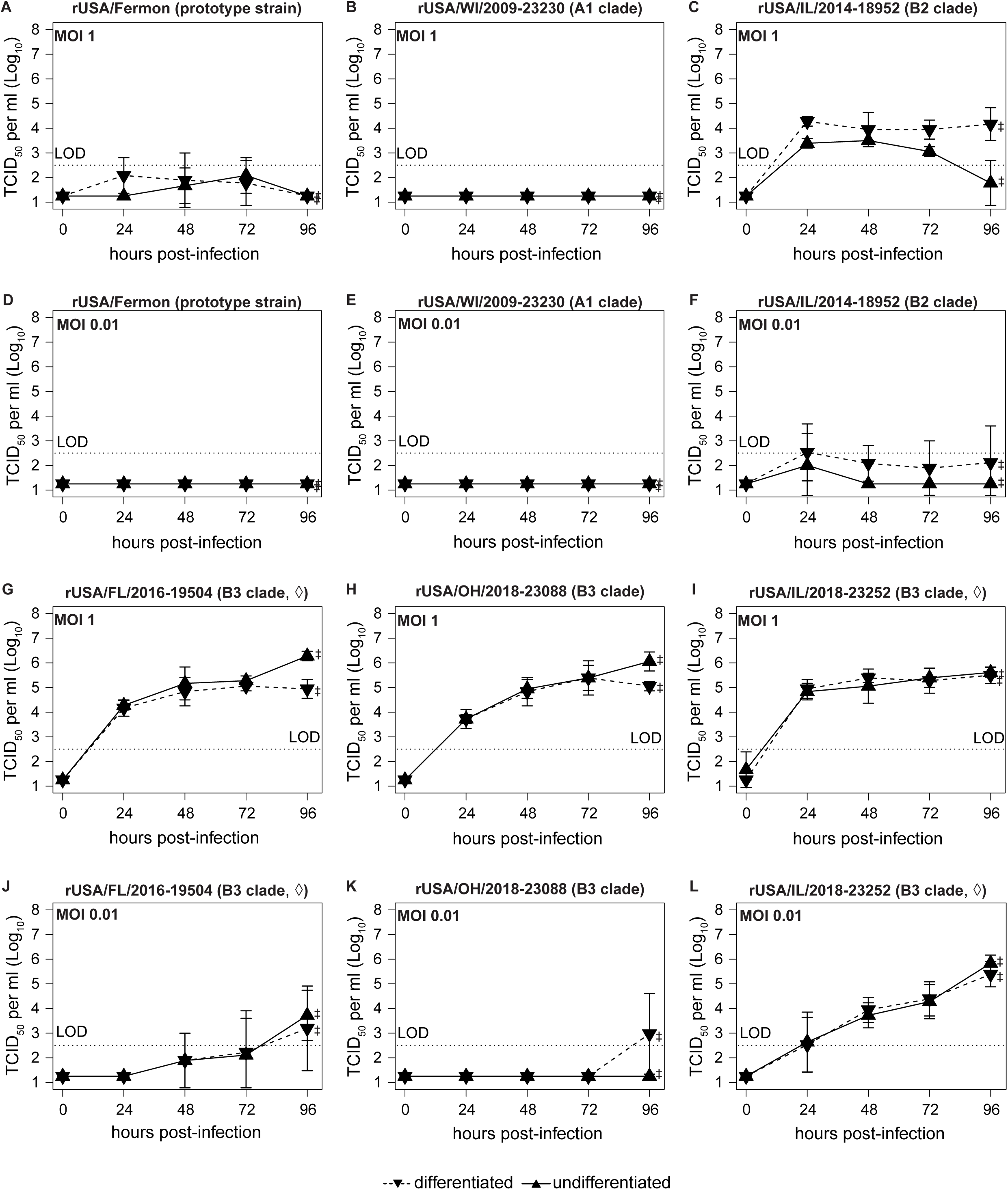
Recombinant B3 clade strains replicate in neurons. SH-SY5Y cells were either differentiated into mature neurons (dashed lines) or maintained in growth medium (solid lines). After 72h, cells were infected in triplicate with the indicated rEV-D68 strains at an MOI of 1 **(A-C** and **G-I)** or 0.01 **(D-F** and **J-L)**. Supernatants were collected at the indicated time points. Endpoint titers represent combined cell lysate and supernatant, as denoted by (‡). (◊) denotes confirmed AFM. Viral titers were quantified by TCID_50_ assay on RD cells. Error bars indicate standard deviation.

Brown *et. al.* reported that differentiation of SH-SY5Y cells into mature neurons is not a requirement for efficient infection or replication of many clinical isolates of EV-D68 in these cells (35). However, we observed differences in growth characteristics in SH- SY5Y cells of some strains, most notably rUSA/IL/2014-18952, that appeared to be dependent on cellular differentiation (**Figure 4, dashed lines**). We quantified this by performing nonlinear regression on viral titers in SH-SY5Y cells at an MOI of 1, where all recombinant B2 and B3 clade strains replicated efficiently (**Figure 4C** and **4G-I**). Using time as the sole predictor of viral titer accurately modeled the majority of titer variance in all strains (**Table 1**). Inclusion of cellular differentiation state had variable impact on the model, with the greatest effect seen in the B2 clade strain rUSA/IL/2014- 18952 (**Figure 4C**). These data are consistent with an interpretation that while rUSA/IL/2014-18952 can infect neurons at any differentiation state, replication is more efficient in differentiated neurons. More modest effects were seen for rUSA/FL/2016- 19504 and rUSA/OH/2018-23088, which both had higher endpoint titers in undifferentiated than differentiated SH-SY5Y cells (**Figure 4G** and **Figure 4H**, respectively). No improvement was seen in the model for rUSA/IL/2018-23252 (**Figure 4I**). Overall, these data suggest that replication of recombinant B3 clade viruses is not restricted by differentiation state.

**Table 1:** Dependence of rEV-D68 viral titer on cellular differentiation state in SH-SY5Y cells. Nonlinear regression was performed on viral titers in undifferentiated or differentiated SH-SY5Y cells. The residual sum-of-squares was used in model selection. Growth kinetics of rEV-D68 were then modeled by a polynomial curve. Time and cellular differentiation state were evaluated as predictors of viral titer for each rEV-D68 strain.

| Strain | Predictor(s) |  |  |  |  |  |
| --- | --- | --- | --- | --- | --- | --- |
|  | Time |  |  | Time and differentiation state |  |  |
|  | Adjusted R<sup>2</sup> | Log likelihood | AIC | Adjusted R<sup>2</sup> | Log likelihood | AIC |
| **rUSA/IL/2014-18952 (B2 clade)** | 0.51 | -36.0 | 80.0 | 0.74 | -25.4 | 62.8 |
| **rUSA/FL/2016-19504 (B3 clade)** | 0.88 | -24.5 | 57.0 | 0.90 | -20.0 | 51.9 |
| **rUSA/OH/2018-23088 (B3 clade)** | 0.93 | -16.5 | 41.0 | 0.94 | -13.5 | 38.9 |
| **rUSA/IL/2018-23252 (B3 clade)** | 0.84 | -27.4 | 62.8 | 0.83 | -27.4 | 66.7 |

### Human spinal cord organoids are susceptible to recombinant B3 clade virus infection

Our studies thus far suggest that recombinant B3 clade strains can infect and replicate in cells of the neuroblastoma SH-SY5Y cell line. However, the spinal cord is comprised of intricate neural circuits with over twenty neuron subclasses (40). We sought to determine whether recombinant B3 clade viruses could establish infection in a more complex model of the spinal cord. Therefore, we further investigated neurotropism of this panel of rEV-D68 strains in three-dimensional spinal cord (3DiSC) organoids. Under 3DiSC differentiation conditions, human spinal cord organoids (hSCOs) are differentiated from human inducible pluripotent stem cells for fourteen days and develop a continuous dorsal neuroepithelial tissue architecture (40). These hSCOs recapitulate more of the cellular heterogeneity found in the spinal cord and we have previously shown that they are susceptible to clinical EV-D68 isolates from contemporary clades, including the B3 clade (36).

We first examined growth kinetics of each rEV-D68 strain in hSCOs. Consistent with our results in SH-SY5Y cells and with clinical isolates (36), control rEV-D68 strains did not produce infectious virus throughout the duration of this experiment (**Figure 5A, red diamonds and squares**). Similarly to what we observed in differentiated SH-SY5Y cells at high MOI, the B2 clade rUSA/IL/2014-18952 strain exhibited sustained replication in hSCOs, with a peak infectious virus titer of 10^5^ TCID_50_ per ml (**Figure 5A, green circles**). Of the B3 clade strains, only rUSA/IL/2018-23252 replicated efficiently in hSCOs, with infectious virus first detected between 24 and 48 hpi and titers continuing to rise throughout the time course (**Figure 5A, inverse purple triangles**). Replication of rUSA/FL/2016-19504 and rUSA/OH/2018-23088 was markedly delayed in hSCOs, in a pattern consistent with that observed in differentiated SH-SY5Y cells at low MOI (**Figure 5A, purple triangles and diamonds, respectively**). Based on these findings, we conclude hSCOs are permissive to recombinant B3 clade strains, but viral replication is more restricted than in SH-SY5Y cells.

**Figure 5.**
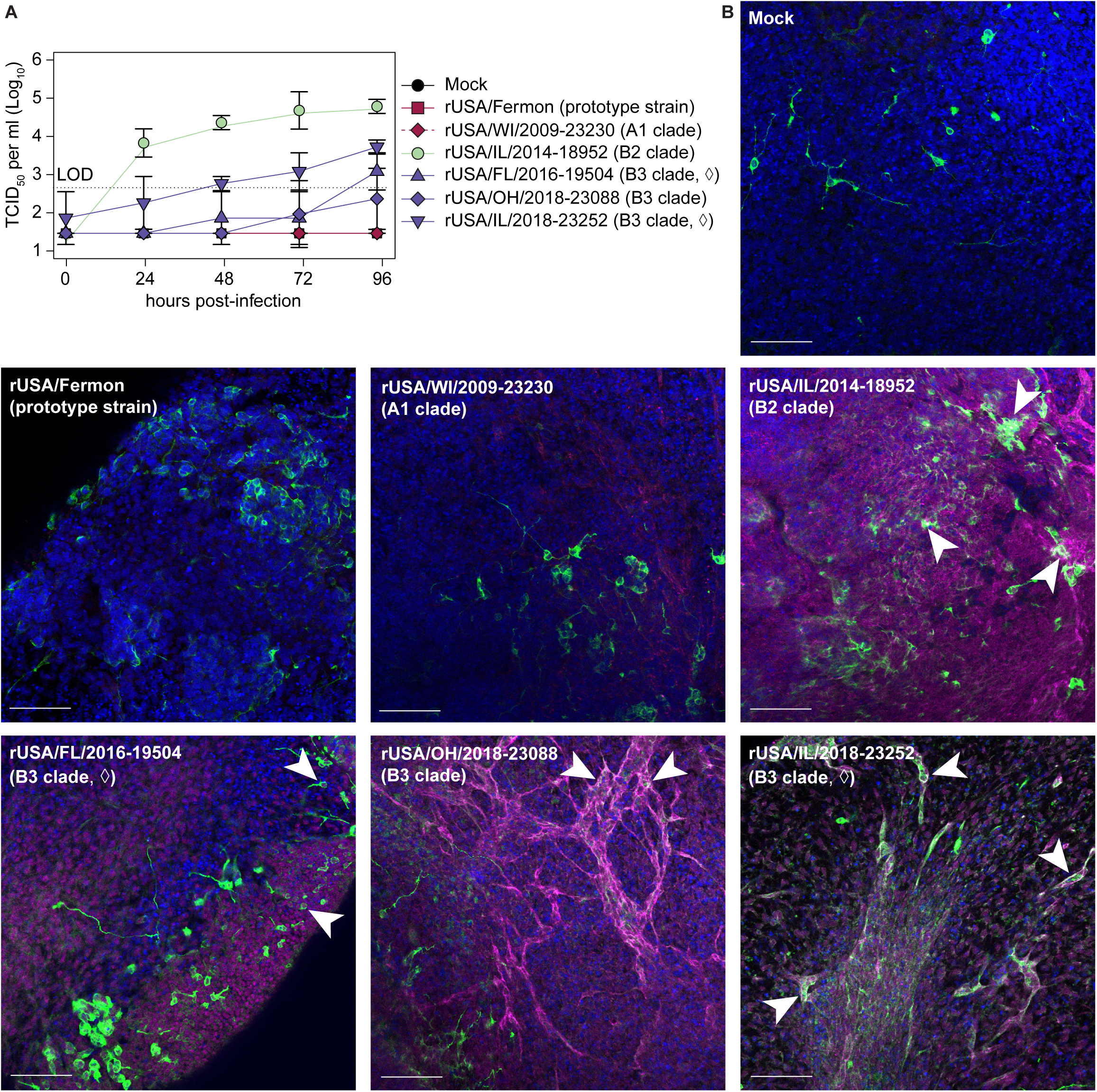
Human spinal cord organoids are susceptible to recombinant B3 clade strain infection. Human inducible pluripotent stem cells were differentiated into 3- dimensional spinal cord (3-DiSC) organoids for 14 days. Pools of 8-12 organoids were either mock-infected or infected with rEV-D68 at 10^6^ PFU per pool in triplicate. (◊) denotes confirmed AFM. **(A)** Viral titer in supernatant was quantified by TCID_50_ assay on RD cells. Error bars indicate standard deviation. **(B)** Organoids were fixed at 96 hpi and stained by immunofluorescence for 4′,6-diamidino-2-phenylindole (DAPI) in blue, a neuronal marker class III β-tubulin (Tuj1) in green, or VP1 in magenta. White arrowheads denote colocalization between VP1 and Tuj1. Scale bars, 100 μm.

To confirm that all B3 clade strains established infection in hSCOs, we additionally performed immunofluorescence staining of hSCOs for VP1, the viral capsid protein. Growth kinetics of rEV-D68 strains in hSCOs mirrored replication kinetics in differentiated SH-SY5Y neuroblastoma cells. Therefore, we examined whether VP1 in infected hSCOs colocalized with class III β-tubulin (Tuj1), an early neuronal differentiation marker (41). We collected hSCOs at 96 hpi, when infectious virus was detected for all recombinant B2 and B3 clade strains and processed for immunofluorescence. Tuj1 staining was readily detected in hSCOs at this time point (**Figure 5B, green**). Minimal to no VP1 staining was observed in hSCOs that were mock-infected or infected with control strains; in contrast, robust VP1 staining was detected in all hSCOs infected with recombinant B2 and B3 clade strains (**Figure 5B, magenta**). Modest VP1 staining was observed for rWI/2009-23230, but no infectious virus was detected for this strain throughout the time course (**Figure 5A, red diamonds**). Conversely, VP1 staining was prominent in hSCOs infected with rUSA/FL/2016-19504 and rUSA/OH/2018-23088, despite the limited replication seen for these strains in hSCOs overall (**Figure 5A, purple triangles and diamonds**). Colocalization of VP1 with Tuj1 was seen in all B2 and B3 clade rEV-D68 strains (**Figure 5B, arrowheads**). Altogether, these data demonstrate that hSCOs are susceptible to recombinant B3 clade virus infection.

## DISCUSSION

The polymerases encoded by RNA viruses lack proofreading enzymes, leading to the replication infidelity and high mutation frequencies characteristic of viruses like EV-D68 (reviewed in (42, 43)). For such viruses, a robust reverse genetics system is not only integral to studies of viral pathogenesis but can greatly improve reproducibility of experimental findings between independent research groups whose lab stocks of clinical isolates may differ. This has contributed to controversial findings for EV-D68, such as whether the prototype strain Fermon is neurotropic (9, 10, 35–38, 44). Yet a comprehensive reverse genetics system for multiple clades of EV-D68 is lacking. Here, we demonstrate reliable recovery of infectious virus from EV-D68 infectious clones by direct transfection of infectious clone and T7 polymerase plasmids into 293T cells, alleviating this limitation on further studies of EV-D68 pathogenesis. While one study found that high titer infectious virus is not readily detected in 293T cells (34), this was not the case in our study. Transfection efficiency played a key role in choice of cell line and likely contributes to this discrepancy. Even after extensive optimization of different commercial transfection reagents and plasmid DNA at a range of dosages, fewer than half of RD cells took up DNA, which undoubtedly impacts DNA delivery and the number of rounds of viral replication required to produce infectious virus. The characteristics of each rEV-D68 strain analyzed in this study are summarized in **Table 2**.

**Table 2:**
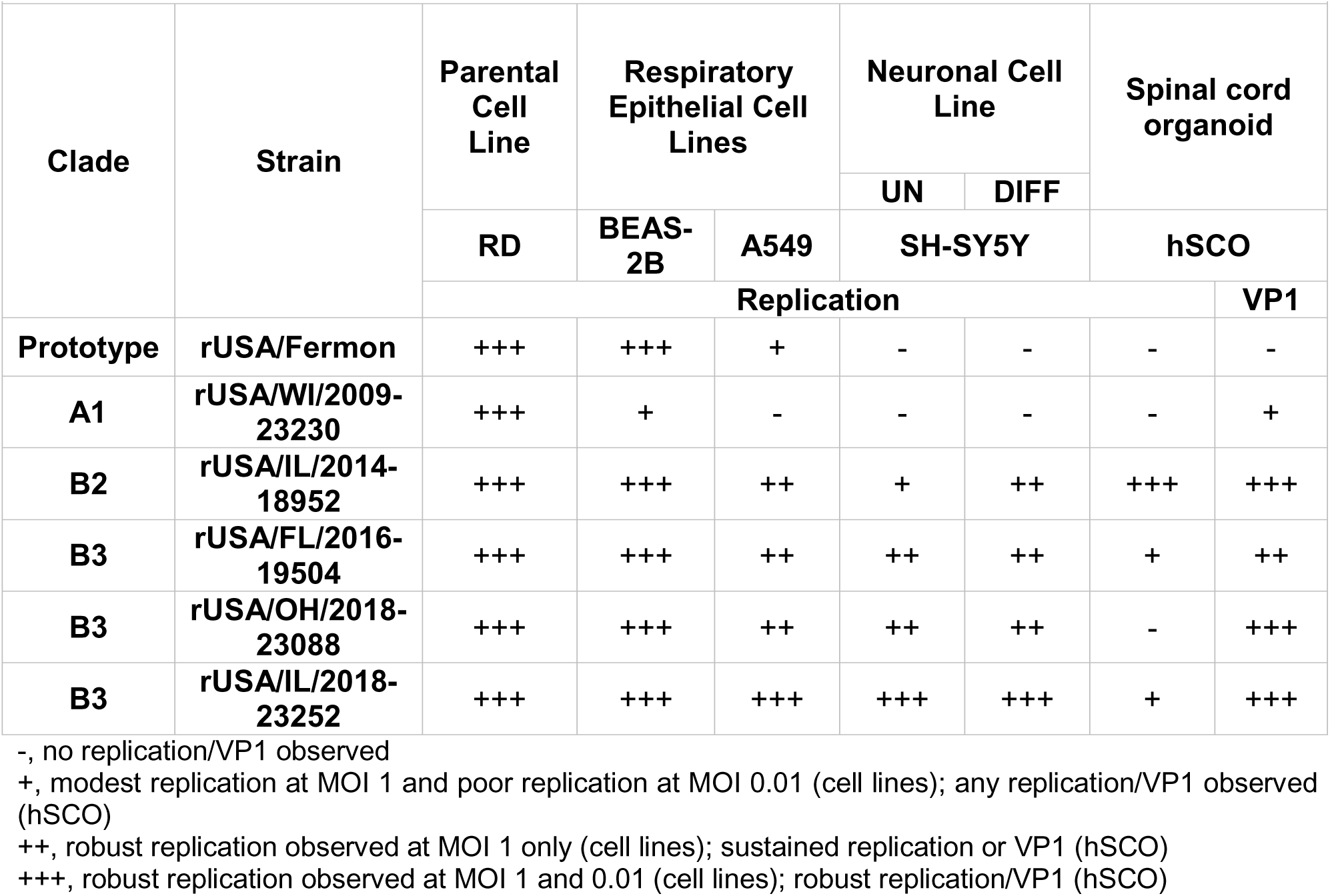
Summary of growth characteristics of rEV-D68 strains. Growth kinetics in human cell lines (RD, BEAS-2B, A549, or SH-SY5Y) or human iPSC-derived spinal cord organoids (hSCO) are summarized under the Replication heading. VP1 expression in hSCOs stained by immunofluorescence are summarized under the VP1 heading. UN, undifferentiated; DIFF, differentiated.

While nearly all rEV-D68 strains replicated efficiently in respiratory epithelial cells, replication was highly constrained in A549 cells compared to BEAS-2B cells. The inverse relationship was found for respiratory syncytial virus (RSV), which more effectively replicates in A549 cells than BEAS-2B cells (45). While both cell lines were initially isolated from human lungs, A549 cells are classified as alveolar type II basal epithelial cells, originally cultured from tissue of an explanted lung adenocarcinoma tumor (46). BEAS-2B cells were instead cultured from autopsy tissue in a non-cancerous normal bronchial epithelium and later immortalized with SV40 T antigen (47). During infection with RSV, A549 cells express high levels of IFN-β-, IFN-λ-, and NF-κB- inducible proinflammatory cytokines, whereas interferon stimulated genes, pattern recognition receptors, and other antiviral genes are upregulated in BEAS-2B cells (45). Given that replication differences between these two cell lines were frequently exacerbated at lower MOI in our study, infection with rEV-D68 likely induces greater antiviral signaling in A549 cells than in BEAS-2B cells. Thus, discrepancies in biological origin, subsequent propagation, and antiviral responses of these cell lines could contribute to the differences observed in viral tropism of rEV-D68.

Recombinant B3 clade viruses were neurotropic in our models, but differed from the paralytogenic B2 clade strain rUSA/IL/2014-18952. Replication of rUSA/IL/2014-18952 was sensitive to cellular differentiation state and MOI in SH-SY5Y cells but was highly efficient in hSCOs, which might suggest that undifferentiated neurons are not a major target for this strain within the human spinal cord. Given that hSCOs are both differentiated and highly heterogeneous (40), replication of this strain may be mediated by differentiated neurons or other non-neuronal cell lineages. Differentiated SH-SY5Y cells release less extracellular virus than undifferentiated SH-SY5Y cells during lytic HSV-1 infection, which is associated with elevated expression of the antiviral protein cyclic GMP-AMP synthase (cGAS) (48). Therefore, differential antiviral responses could be responsible for the abortive phenotype observed for rUSA/IL/2014-18952 in undifferentiated SH-SY5Y cells. In contrast, the B3 clade rUSA/IL/2018-23252 strain replicates efficiently in both SH-SY5Y and hSCO models. This strain was isolated in 2018 from a nasopharyngeal swab of a patient with confirmed AFM in 2018, a year when there were the greatest number of US cases of AFM (18, 22). This strain had the broadest cellular tropism overall, frequently replicating to the highest titers of all strains independent of cell type or MOI, including in A549s, which were least permissive to rEV- D68. Taken together, these data could reflect adaptations rendering this virus more efficient at evading antiviral signaling pathways in diverse cell types. Ultimately, the ability to recover infectious virus from diverse rEV-D68 strains has provided exciting new avenues for investigation into novel viral determinants of AFM.

## MATERIALS AND METHODS

### Cells

Human embryonic kidney (HEK) 293T cells (ATCC CRL-3216), human adenocarcinoma lung epithelial A549 cells (ATCC CCL-185), human rhabdomyosarcoma (RD) cells (ATCC CCL-136), human lung bronchial epithelial BEAS-2B cells (ATCC CRL-3588), and human neuroblastoma SH-SY5Y cells (ATCC CRL-2266) were obtained from ATCC. Baby hamster kidney (BHK)-21 cell clone BSR-T7/5 has been previously described (32). Human iPSCs were obtained from STEMCELL Technologies (SCTi003- A). A549, 293T, RD and BEAS-2B cells were maintained in Dulbecco’s Modified Eagle’s Medium (DMEM, Corning) supplemented with 10% fetal bovine serum (FBS, Biowest) and 1% penicillin/streptomycin (Gibco). BSR-T7 cells were maintained in Eagle’s Minimum Essential Medium (EMEM, Gibco) supplemented with 10% FBS, 1% sodium pyruvate (Corning), 1% nonessential amino acids (Corning), and 1% penicillin/streptomycin (Corning). SH-SY5Y cells were maintained in EMEM supplemented 1:1 with Ham’s F12 nutrient mix (Gibco), 10% FBS, 1% sodium pyruvate, 1% nonessential amino acids, and 1% penicillin/streptomycin. Where specified, SH- SY5Y cells were differentiated into mature neurons by the reduction of the FBS concentration in the culture medium to 3% and the addition of 100 nM retinoic acid (Tocris) for 72h, as previously described (35). Human iPSCs were maintained in mTeSR^TM^ Plus medium (STEMCELL Technologies) in flasks coated with 150 µg/mL Cultrex (R&D Systems). All cells were routinely tested for mycoplasma contamination with a PCR-based detection kit either in-house (Sigma) or at the University of Arizona Genetics Core (Applied Biological Materials Inc.). Cells were maintained in sterile cell culture incubators at 37°C, 5% CO_2_ unless otherwise stated. Low-passage cultures were used to ensure purity of the culture.

### Cell line authentication

All cell lines used in this study were authenticated by short tandem repeat (STR) profiling. Genomic DNA was extracted from cells using the DNeasy Blood and Tissue Kit (Qiagen) following the manufacturer’s guidelines and shipped to the University of Arizona Genetics Core for STR profiling. Briefly, genomic DNA was genotyped for 15 autosomal STR loci and amelogenin (X/Y) with the Powerplex 16HS PCR kit (Promega). Positive control (gDNA 2800M, Promega) and negative control (water) were also amplified to confirm the accuracy of allelic calls and that reactions were free of contaminating genetic material. PCR products were separated by capillary electrophoresis using an AB 3730 DNA Analyzer. Applied Biosystems Internal Size Standard GeneScan500-LIZ was used to standardize allele size calls for each sample. Samples were run on a 36 cm capillary array (Applied Biosystems). Electropherograms were analyzed and allelic values assigned using Soft Genetics, Gene Marker Software Version 3.0.1. Alleles were matched to the STR profile recorded in the Deutsche Sammlung von Mikroorganismen und Zellkulturen (DSMZ) Leibniz Institute database (when reference profile is available). An American National Standards Institute (ANSI) standard of a minimum 80% match threshold indicates a shared genetic history.

### Plasmids and transfection optimization

The following EV-D68 infectious clones were obtained through BEI Resources, National Institute of Allergy and Infectious Diseases (NIAID), National Institutes of Health (NIH): rUSA/Fermon was rescued from pUC19-R-Fermon (NR-52375), rUSA/WI/2009-23230 from pUC-R23230 (NR-52377), rUSA/IL/2014-18952 from pUC19-EVD68_49131 (NR- 52011), rUSA/OH/2018-23088 from pUC-R23088 (NR-52379), rUSA/IL/2018-23252 from pUC-R23252 (NR-52380), and rUSA/FL/2016-19504 from pUC-R19504 (NR- 52378).

Sequences of plasmid stocks of infectious clones were verified by whole-plasmid, long-read sequencing (Oxford Nanopore, Plasmidsaurus). The following discrepancies were found between infectious clone sequences and their GenBank counterparts: pUC- R19504 and pUC-R23088, 2A S11F; pUC-R23088 and pUC-R23252, 3D D43N. The pCAGGS T7 opt plasmid was obtained through Addgene (#65974).

Transfection conditions were empirically determined for RD, BSR-T7, and 293T cells. Briefly, each cell type was grown in tissue culture plates. Cells were transfected with a pcDNA 3.1 (+) CT-GFP reporter construct (Invitrogen) and either TransIT-LT1 transfection reagent (Mirus Bio) or Lipofectamine 3000 (Invitrogen) at a range of concentrations according to the manufacturer’s specifications. GFP expression was monitored at 24-72 hours post-transfection using an EVOS FL Auto 2 cell imaging system (Invitrogen). Transfection conditions yielding maximal GFP expression and minimal toxicity were selected and applied to virus rescue. Lipofectamine 3000 was selected for RD cells and TransIT-LT1 transfection reagent was chosen for 293T and BSR-T7 cells.

### Virus rescue optimization

Virus rescue conditions were empirically determined for RD, BSR-T7, and 293T cells. Briefly, each cell type was grown in tissue culture dishes. Transfection conditions for each cell type were scaled up from the GFP transfection experiment and applied to pUC19-EVD68_49131 to rescue rUSA/IL/2014-18952. RD and 293T cells additionally received pCAGGS T7 opt expression construct at the following ratios to infectious clone: 10:1, 5:1, 1:1, 1:5, or 1:10. Mock-transfected cells received transfection mixture without DNA. Transfections were performed according to manufacturer specifications and cells were incubated at 33°C, 5% CO_2_ until collection. Plaque assays were performed on P0 stocks of rUSA/IL/2014-18952 and rescue conditions were selected based on these titers.

The full panel of recombinant viruses, including a fresh stock of rUSA/IL/2014-18952, was rescued from infectious clones in 293T cells. Cells were grown on tissue culture dishes. Transfection mixtures were supplied with pCAGGS T7 opt expression construct and the indicated infectious clone at a ratio of 10:1. Transfections were performed with TransIT-LT1 transfection reagent (Mirus Bio) according to manufacturer specifications. Mock-transfected cells received transfection mixture without DNA. Cells were incubated at 33°C, 5% CO_2_ until collection.

P0 stocks were amplified one additional time on RD cells. Briefly, RD cells were grown on tissue culture dishes. Half of the P0 stock was added to cell monolayers and cells were rocked at room temperature for 1 hour to adsorb virus. Mock-infected cells received cell culture media. Inocula were aspirated from dishes and cell growth medium was replenished. Cells were incubated at 33°C, 5% CO_2_ until collection. P1 stocks were harvested and clarified by ultracentrifugation at 12,000 rpm for 10 minutes at 16°C. Viral stocks were stored at −80°C.

### Phylogenetics

The following GenBank accession numbers were used to source VP1 sequences of each recombinant strain in this study: NC_038308 (USA/Fermon), MN240506 (USA/WI/2009-23230), KM851230 (USA/IL/2014-18952), MN245982 (USA/OH/2018- 23088), MN246015 (USA/IL/2018-23252), and KX675261 (USA/FL/2016-19504). Sequences were read into R (version 4.3.2) and analyzed with the DECIPHER (version 2.30.0), ape (version 5.7-1), and phangorn (version 2.11.1) packages. A maximum-likelihood tree was reconstructed from a multiple sequence alignment and rooted to the prototype strain (USA/Fermon). A general time reversible model was selected based on log likelihood and Akaike information criterion (AIC). Bootstrapping was performed with 1,000 trees.

### Growth kinetics in human cell lines

RD, A549, BEAS-2B, and SH-SY5Y cells were grown in tissue culture plates. Half of the SH-SY5Y cells were differentiated into mature neurons. Recombinant EV-D68 was diluted to an MOI of 1 or 0.01 in PBS containing calcium and magnesium (Corning). Cells were inoculated with the indicated strain or mock-infected with diluent. Inocula were adsorbed onto cells at room temperature for 1 hour with rocking. After 1 hour, the inoculum was aspirated, cells were washed three times with PBS without calcium/magnesium (HyClone), and growth medium was added to cells. An aliquot was taken from each well and designated 0 hpi. Cells were incubated at 33°C, 5% CO_2_ for the course of infection, with supernatants collected at 24, 48, 72, and 96 hpi. At the endpoint of the assay, cells were scraped and harvested with their respective supernatants. Samples were stored at −80°C.

### Differentiation & propagation of human spinal cord organoids

3-DiSC hSCOs were differentiated from iPSC cells as previously described (36). Briefly, iPSC cells were dissociated into 96-well round-bottom low-adhesion plates using Accumax (Sigma). Cells were seeded at a density of 9,000 cells per well. Cells were plated in differentiation medium for 14 days, with media replenished every 3 days. After 14 days, organoids were transferred to a tissue culture plate and cultured in suspension in N2B27 medium [DMEM/F-12 (Gibco), neurobasal medium (Gibco) (1:1), 0.5% (vol/vol) N2 supplement (ThermoFisher) and 1% (vol/vol) B27 supplement without vitamin A (Gibco)] supplemented with 1 mM L-glutamine (ThermoFisher), 0.1 mM β- mercaptoethanol, 0.5 µM ascorbic acid, 10 ng/mL brain-derived neurotrophic factor (BDNF, STEMCELL Technologies), 10 ng/mL glial cell line-derived neurotrophic factor (GDNF, STEMCELLL Technologies), and 100 nM retinoic acid (Tocris).

### Growth kinetics in human spinal cord organoids

3-DiSC hSCOs were inoculated with rEV-D68 at 14 days post-propagation in pools of 8-12. Viruses were diluted to 10^5^ PFU per pool in N2B27 medium. Pools were either infected with the indicated strain or a mock inoculum of PBS. Virus was adsorbed on hSCOs for 1 hour at room temperature with rocking. After 1 hour, inoculum was aspirated and organoids were washed 3 times with PBS without calcium/magnesium (HyClone). Pools were transferred to new plastic to ensure there was no residual inoculum present and fresh growth medium was added. hSCOs were incubated at 33°C, 5% CO_2_ for the duration of the experiment. Supernatants were taken at 0, 24, 48, 72, and 96 hpi and stored at −80°C. At 96 hpi, hSCO were fixed with 4% paraformaldehyde in PBS (ThermoFisher) for immunofluorescent staining as indicated.

### Viral titers

Infectious virus was quantified either by standard plaque assay or 50% tissue culture infectious dose assay (TCID_50_) on RD cells, as indicated. TCID_50_ titers were enumerated by the Spearman-Kärber method (49).

### Immunofluorescence staining

Immunofluorescence staining on fixed hSCOs was completed as previously described (50). Briefly, hSCOs were washed with 1% PBS-BSA and PBS-T at 4°C. The remaining washes and antibody dilutions were in organoid wash buffer (0.1% Triton X-100 (v/v), 0.2% BSA (m/v) in PBS). Primary antibodies VP1 (Genetex, GTX132313) and Tuj-1 (ThermoFisher, 480011) were used at a 1:250 dilution. Secondary antibodies Alexa Fluor^TM^ 647 (Thermo Fisher, A-21245) and Alexa Fluor^TM^ 488 (A-11001) were used at a 1:1000 dilution. DAPI (4’,6-diamidino-2-phenylindole, dihydrochloride) (ThermoFisher, D1306) was used at a 1:300 dilution. hSCOs were cleared for 30 minutes with 60% glycerol in 2.5 M fructose before mounting.

### Confocal imaging

Microscope slides were imaged on a Leica Stellaris 5 confocal microscope equipped with a pulsed white light laser as an excitation source and an acousto-optical beam splitter and Leica Hybrid Detectors. Volumetric Z-stacks were imaged with a 20X objective. Sequential scanning was used with 2X line averaging by frame. The following parameters were used: dsDNA was measured 403 nm, Tuj1 was measured at 488 nm, and VP1 was measured at 647 nm. Images were processed with FIJI 2 version 2.9.0.

### Statistical analysis

Linear regression was performed in R (version 4.4.1) to analyze the relationship between the ratio of T7 polymerase to infectious clone and viral titer. Nonlinear regression was performed on viral titers in SH-SY5Y cells. Growth kinetics of rEV-D68 in SH-SY5Y cells were modeled by a polynomial curve. This model was selected based on the residual sum-of-squares of several potential curves for each rEV-D68 strain. Time and cellular differentiation state were evaluated as potential predictors of viral titer for each rEV-D68 strain. Model parameters were supported or rejected based on the adjusted R^2^, log likelihood, and AIC.

## ACKNOWLEDGMENTS

This work is funded by the Richard King Mellon Foundation (M.C.F.) and M.C.F. is supported by NIH NIAID K08 AI171177. The funders had no role in study design, data collection and interpretation, or the decision to submit the work for publication. Confocal images were obtained in the Cell Imaging Core Facility at the UPMC Children’s Hospital of Pittsburgh Rangos Research Center. We thank John V. Williams for his review of the manuscript and Gabrielle Aguglia for her laboratory support. We thank members of the Williams, Freeman, Dermody and Dolan labs for helpful discussions in the development of this manuscript.

